# Methods for imaging intracellular calcium signals in the mouse mammary epithelium in 2- and 3-dimensions

**DOI:** 10.1101/2024.02.27.582265

**Authors:** Mathilde Folacci, Silke B. Chalmers, Felicity M. Davis

## Abstract

The mammary gland has a central role in optimal mammalian development and survival. Contractions of smooth muscle-like basal (or myoepithelial) cells in the functionally mature mammary gland in response to oxytocin are essential for milk ejection and are tightly regulated by intracellular calcium (Ca^2+^). Using mice expressing a genetically encoded Ca^2+^ indicator (GCaMP6f), we present in this chapter a method to visualise at high spatiotemporal resolution changes in intracellular Ca^2+^ in mammary epithelial cells, both *in vitro* (2D) and *ex vivo* (3D). The procedure to optimally prepare mammary tissue and primary cells is presented in detail.

## 1.0 Introduction

Essential and eponymous, the mammary gland fosters optimal development and survival at the most vulnerable stage of mammalian life [1–3]. In mice, the post-pubertal mammary gland consists of a network of branching ducts that are situated within a rich, vascularised stromal matrix containing fibroblasts, adipocytes, and immune cells [4, 5]. During gestation and early lactation, the mammary epithelium undergoes alveologenesis, a process that involves the rapid expansion and reorganisation of the epithelium, enabling it to synthesise, store, and secrete milk [6]. Alveolar structures created during this process consist of an inner layer of milk synthesising luminal cells encaged by star-shaped, contractile basal cells [6, 7]. Upon weaning, the gland undergoes a highly coordinated phase of regression—known as post-lactational involution—enabling it to return to a near pre-pregnant state, ready for future recurrent cycles of expansion and milk production with subsequent pregnancies [8].

The role and function of the mammary gland is inextricably linked to Ca^2+^ [9, 10]. Historically considered simply a nutritional constituent of milk, Ca^2+^ is now understood to be a key regulator of various mammary functions through its role as a ubiquitous second messenger [9, 11–14]. Ca^2+^ influx via ORAI1 is required for oxytocin-induced contractions of basal cells [15, 16]. Moreover, the onset of involution involves a rapid increase in luminal cytosolic Ca^2+^ concentrations, likely due to the early downregulation of apically-expressed plasma membrane Ca^2+^ ATPase (PMCA)2 [17].

Recent improvements in our understanding of Ca^2+^-mediated control of mammary biology largely come from advances in live imaging and Ca^2+^ signalling fields. Improved methods for visualising the mammary epithelium have enabled researchers to explore mammary morphogenesis and function, from *in vitro* to *in vivo* scales (**Figure 1**) [15, 16, 18–22]. Concomitantly, transgenic mouse models enabling targeted expression of fast and sensitive genetically encoded Ca^2+^ indicators (GECIs) have become widely available [4, 16, 23]. Together, these advances enable researchers to probe mammary Ca^2+^ signalling with a hitherto unknown clarity. Herein, we describe methods routinely used within our laboratory to visualise mammary Ca^2+^ signalling across 2D and 3D scales, while offering insights and suggestions regarding choice of experimental technique and mouse model.

**Figure 1:**
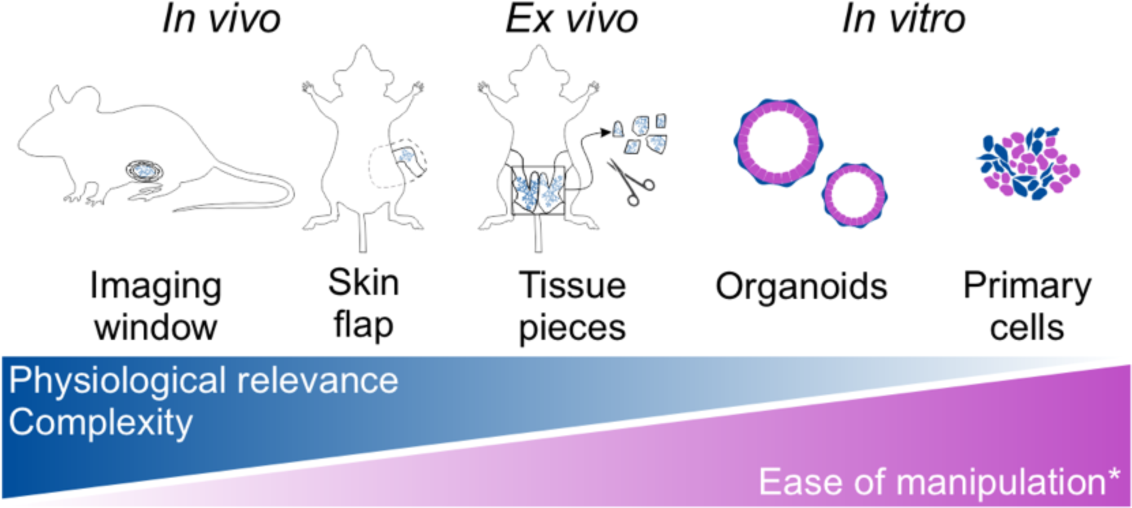
Methods to image the mouse mammary gland range from relatively simple 2D cell preparations to serial *in vivo* manipulations. Complexity (time, expertise, equipment and analysis pipelines) tends to increase with 3D/4D and *in vivo* methods; however, these methods offer the advantage of improved physiological relevance. Asterisk signifies that not all methods are currently available or readily reproducible for all stages of mammary gland development, e.g., while it is routine to isolate healthy primary cells from non-pregnant mammary tissue, it is more difficult isolate healthy cells from lactating mouse mammary tissue. Similarly, methods for implanting imaging windows before or during gestation remain a major challenge for intravital imaging during alveologenesis and lactation.

### Model considerations

Model selection, including choice of GECI and the method of cell type specific targeting, influences suitability for flow-on applications. Of the range of GECIs that have been expressed in mouse models, only GCaMP6f has been assessed in mammary epithelia to date [16, 18, 23, 24]. GCaMP6f has the advantages of high sensitivity and fast kinetics; however, as a green, non-ratiometric sensor that exhibits low fluorescence in a Ca^2+^-unbound state, it poses challenges for imaging Ca^2+^ signals in thick, contractile and auto-fluorescent mammary tissue [23, 25]. Previous studies have addressed some of these issues by employing more complex breeding strategies to generate mice expressing both GCaMP6f and a bright, Ca^2+^-insensitive red fluorescent protein (TdTomato) for ratioing [16, 24]. Recently, mouse models with dual GCaMP6f-TdTomato transgene insertion (known as ‘Salsa6f’) have become available through worldwide distributors, ameliorating the time and cost required to obtain both GCaMP6f and TdTomato on the Rosa26 locus purely by breeding [24]. In any case, fluctuations in intracellular Ca^2+^ in GCaMP6f-expressing models should be interpreted as response-over-baseline (i.e. f/f_0_ or Δf/f_0_).

The Cre-driver used to direct expression of the GECI to the mammary cell type of interest is another important consideration in the experimental design [6, 15, 26]. Commonly utilised mammary Cre models include K5-CreERT2, K8-CreERT2, whey acidic protein (WAP) Cre, and beta-lactoglobulin (BLG or LGB) Cre [4, 26–28]. However, these models are not without pitfalls. Use of tamoxifen in creERT2 models can affect gland development and may also increase the likelihood of breeding complications (sterility, dystocia, cannibalisation) in breeding studies. Moreover, WAP-Cre and BLG-Cre models require a cycle of pregnancy and lactation for optimal transgene expression, making them time intensive and unsuitable for assessment of Ca^2+^ signalling in the nulliparous state [4, 29]. Whether cell type specific expression is required, however, may depend on the downstream application. The method of primary cell isolation discussed in this paper produces a mixed population of cells; thus, non-specific expression of a GECI would make it difficult to discern cell lineages during live Ca^2+^ imaging experiments [18]. Conversely, the differing morphologies of luminal and basal cells in gestation and lactation may make *in vivo* or *ex vivo* imaging of tissue with non-specific GECI expression viable. Interpretation and analysis of results from these experiments, however, could be challenging [30–32].

Of the methodical approaches depicted in **Figure 1**, all but serial intravital imaging have been employed for mammary Ca^2+^ signalling studies [16, 18]. Each method has its advantages and difficulties, but perhaps more notably, they are also distinguished by which developmental stage they are best suited to (**Figure 1**). This paper will focus on the two most commonly used methods within our laboratory; *ex vivo* imaging of live, thick tissue during gestation or lactation, and *in vitro* imaging of isolated primary mammary epithelial cells from non-pregnant/non-lactating mice. These two methods provide scope for imaging different stages of the mature mammary gland. Of note, many published mammary dissection protocols describe methods that ensure optimal dissection of the abdominal (4^th^) mammary glands, but not the inguinal (5^th^) glands [33–35]. In our experience, the pair of inguinal mammary glands provide valuable tissue for both *ex vivo* imaging and primary cell isolation due to the high ratio of mammary epithelial cells in comparison to surrounding stroma. We therefore refer readers to our previous technical paper for a detailed protocol of mammary gland dissection [4].

### Advantages and disadvantages of *ex vivo* mammary imaging

Tissues isolated during gestation and in lactation are well suited to *ex vivo* imaging (**Figure 2**). At late gestation, mammary alveoli have not yet undergone secretory activation; their smaller diameter means that entire lobuloalveolar units and the subtending duct can be visualised in a single field of view with a 20-30× confocal objective [16]. Lactating tissue, in comparison, contains larger, more mature structures [16]. In non-pregnant mice, most of the ductal epithelium is buried within an adipocyte-rich stroma, making it difficult to visualise mammary ducts and end buds using conventional confocal microscopes.

**Figure 2:**
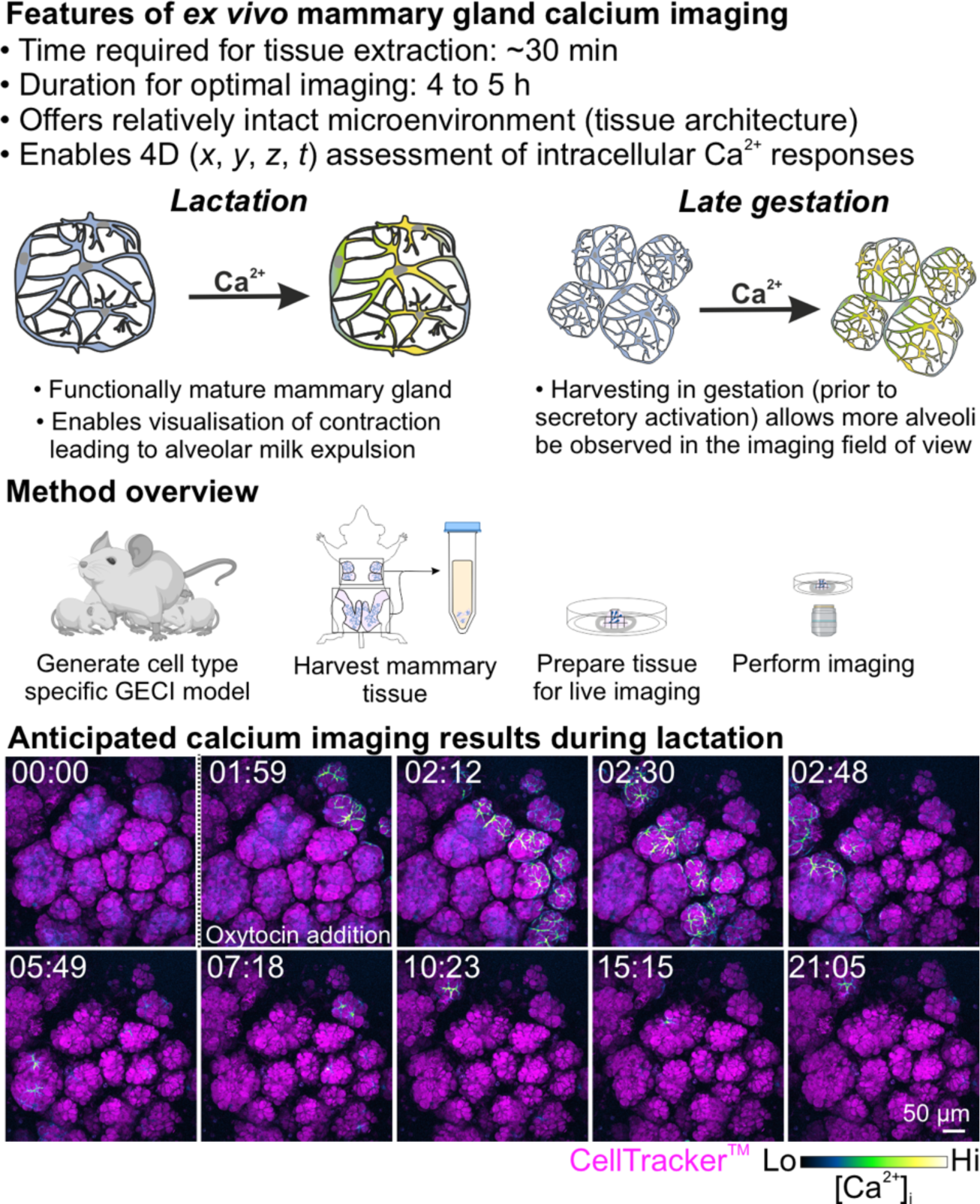
Overview of *ex vivo* Ca^2+^ imaging of basal cells in the mammary gland. Figure shows features of *ex vivo* imaging during lactation and late gestation; schematic representation of the method; and anticipated results (lactation). For the live tissue imaging, time is indicated at the top left (min:sec). Luminal cells are stained with CellTracker (magenta) and changes in cytosolic Ca^2+^ in basal cells expressing GCaMP6f are shown in blue/green/yellow intensity LUT.

One major limitation of *ex vivo* imaging is the limited diffusion of pharmacological agents through thick tissue to the cell type of interest. As will be described below, this issue can be in-part circumvented through delicate dissection of mammary tissue into small pieces, and selection of imaging regions near the edge of the tissue piece. Where relevant, *in vitro* or *in vivo* applications may overcome this challenge and could be considered.

### Advantages and disadvantages of *in vitro* imaging of primary mammary epithelial cells

Isolated primary mammary epithelial cells are a valuable resource for visualising Ca^2+^ signalling in luminal and basal cell populations (**Figure 3**) [18]. In our experience, isolation from non-pregnant/non-lactating mammary tissue provides the highest yield of viable cells, in comparison to tissue isolated during gestation or lactation. Due to the propensity of primary cells to dedifferentiate when cultured on stiff surfaces, we consider isolated mammary cells most suitable for Ca^2+^ imaging up to 24 h post-dissection [18, 36, 37].

**Figure 3:**
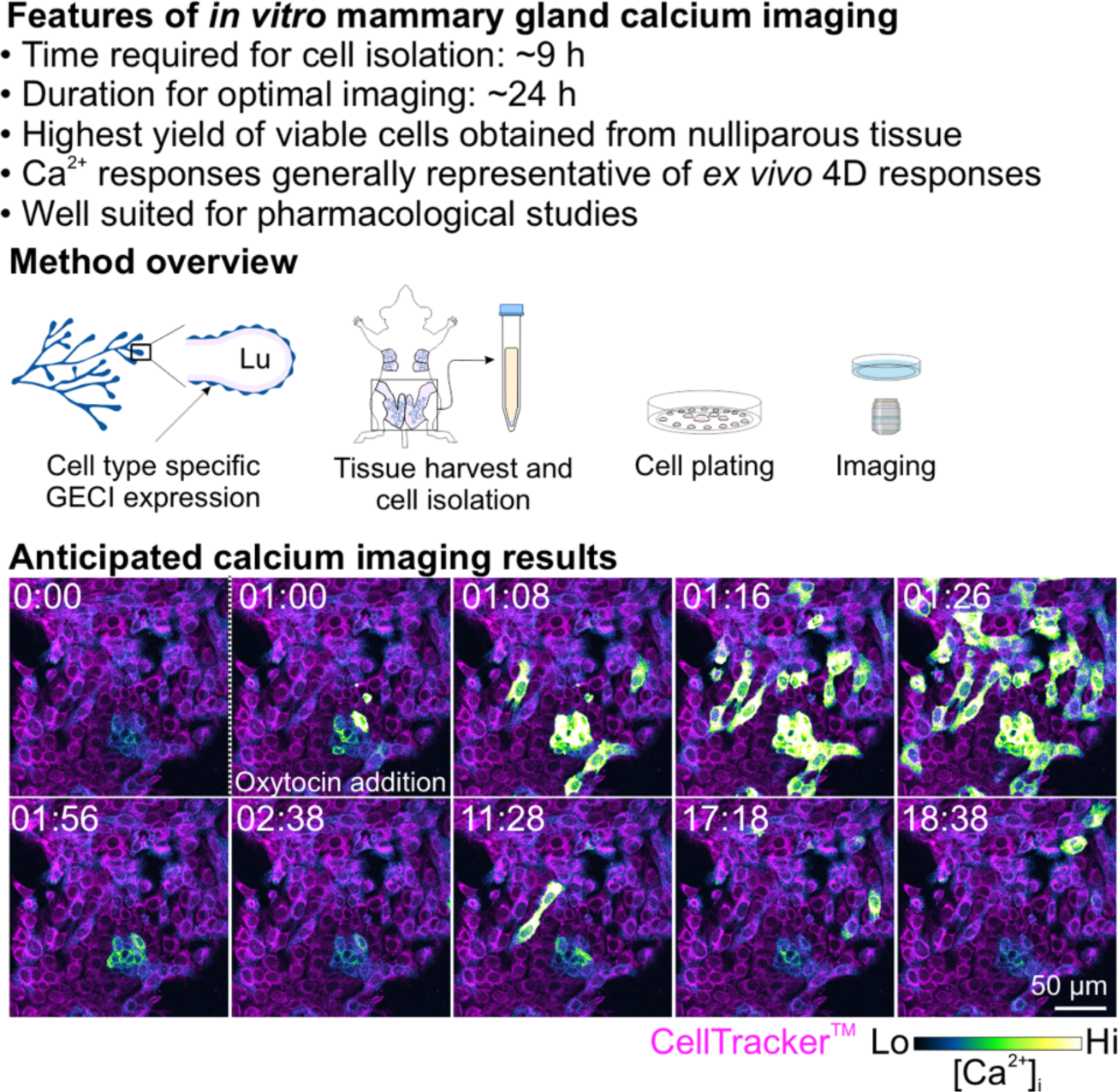
Overview of *in vitro* Ca^2+^ imaging of primary mammary epithelial cells. Isolation of mammary epithelial cells from nulliparous tissue produces monolayers of cells that remain viable for Ca^2+^ imaging studies up to 24 hours post-dissection. The Ca^2+^ signalling responses of these cells are generally representative of responses in *ex vivo* tissue. For the live cell imaging experiment, time is indicated at the top left (min:sec). Cells have been stained with CellTracker (magenta) and changes in cytosolic Ca^2+^ in basal cells expressing GCaMP6f are shown in blue/green/yellow intensity LUT. Lu: lumen.

The cell isolation method described in this chapter utilises the proprietary gentleMACS^TM^ dissociator. The dissociation step can, however, be performed using other published protocols [38–40]. Whilst outside the scope of this chapter, following isolation, primary epithelial cells can additionally be utilised for the development of mammary organoids, which our group has shown are also suitable for Ca^2+^ imaging studies. For further information on this, we recommend previous descriptions of this process [18, 35].

## 2. Materials

Use ultrapure deionized water for all solutions. Storage temperatures are at room temperature unless otherwise specified. Safety measures, ethical approvals, and waste disposal should be followed according to local and national regulations. Materials related to the mouse mating procedure and mammary gland harvest are described in Stewart & Davis [4].

### 2.1 Materials for mammary gland *ex vivo* Ca^2+^ imaging

1. Transgenic mouse expressing a GECI in mammary tissue (**Note 1**)
2. Calcium chloride dihydrate (CaCl_2_.2H_2_O)
3. Magnesium chloride (MgCl_2_) solution (1 M)
4. Sodium phosphate monobasic (NaH_2_PO_4_)
5. Potassium chloride (KCl)
6. 4-(2-hydroxyethyl)piperazine-1-ethane-sulfonic acid (HEPES, C_8_H_18_N_2_O_4_S)
7. Sodium bicarbonate (NaHCO_3_)
8. Sterile disposable 0.2 μm filtration unit (500 mL) and vacuum / vacuum pump
9. Magnetic stirrer and sterile laboratory glassware
10. Sodium chloride (NaCl)
11. D-(+)-Glucose (C<u>_6_</u>H_12_O_6_)
12. Ethanol (70%)
13. Humidified cell culture incubator (37°C, 5% CO_2_)
14. Biosafety cabinet
15. Disposable scalpel blades (#20)
16. Petri dish (140 mm)
17. Centrifuge tube (50 mL)
18. Dulbecco’s Modified Eagle Medium/Nutrient Mixture F-12 (DMEM/F-12) with HEPES and L-glutamine
19. Fetal bovine serum (FBS)
20. Optical grade (0.17 mm, #1.5) glass bottom culture dishes
21. Net and stainless-steel washer as shown in **Figure 4g**, or equivalent.
22. *Optional,* CellTracker Red CMTPX (#C34552 ThermoFisher) (**Note 2**)
23. *Optional,* dimethyl sulfoxide (DMSO) for reconstituting CellTracker Red
24. Oxytocin (#O3251, Sigma Aldrich)
25. CaCl_2_ solution (2 M): Weigh CaCl_2_.2H_2_O and dissolve in H_2_O. Store at 4°C.
26. NaH_2_PO_4_ solution (1 M): Weigh NaH_2_PO_4_ and dissolve in H_2_O. Store at 4°C.
27. HEPES stock solution (10×). For 500 mL: KCl (2.199 g), 1 M MgCl_2_ solution (7 mL), HEPES (11.915 g), 1 M NaH_2_PO_4_ solution (6 mL), NaHCO_3_ (2.1003 g). Weigh and add dry components to the clean glass bottle. Dissolve in H_2_O. Add MgCl_2_ and NaH_2_PO_4_ solutions. Adjust the volume to 500 mL with H_2_O. Pass through a 0.2 μm filtration unit. Store and 4 °C. Maintain sterility and discard after 6 months.
28. Physiological salt solution with 1.8 mM Ca^2+^ (PSS-Ca^2+^). For 500 mL: 10× HEPES stock (50 mL), NaCl (4.0908 g), D-(+)-Glucose (1.0359 g), 2 M CaCl_2_ solution (450 μL). Weigh and add dry components to the clean glass bottle. Dissolve in H_2_O. Add 10× HEPES and 2 M CaCl_2_ solutions. Adjust the volume to approx. 480 mL with H_2_O. Adjust the pH to 7.2-7.3 using NaOH. Make up the final volume to 500 mL with H_2_O. Store at 4 °C and discard after 1 month.
29. Sterile live imaging media: DMEM/F-12 supplemented with 10% FBS (store at 4 °C).
30. Oxytocin stock solution: 1 mM, reconstituted in H_2_O. Aliquot and store at −20 °C.
31. Oxytocin 2× solution: 150 nM, diluted in PSS-Ca^2+^. Prepare fresh on the day of the experiment. Keep on ice until ready to use.
32. *Optional,* CellTracker Red stock solution: 6 mM, reconstituted in DMSO. Store at −20 °C. Avoid excessive freeze-thawing.
33. *Optional,* CellTracker Red working solution: 1.5 µM, diluted from 6 mM stock in live imaging media (subheading 2.1 item 29) and mixed well by vortexing. Prepared on the day of the experiment. Warmed to 37 °C 20 min prior to use.
34. Confocal microscope, preferentially with a resonant scanner.
35. ImageJ software (v1.52e, National Institutes of Health)

### 2.2. Materials for primary mammary cell isolation and *in vitro* Ca^2+^ imaging

1. Female mice (aged 8-34 weeks) expressing GECI of choice in cell type of interest (**Notes 3 and 4**)
2. DMEM/F-12 with HEPES and L-glutamine
3. DMEM, high glucose, without L-glutamine or HEPES
4. 100× GlutaMax^TM^
5. 100× Penicillin/streptomycin: 10,000 units of penicillin, 10,000 μg of streptomycin
6. Leibovitz’s L-15 medium
7. FBS
8. Trypsin from bovine pancreas
9. Collagenase A from *Clostridium histolyticum*
10. Red blood cell lysis buffer (Sigma Aldrich)
11. Dissociation media: Leibovitz’s L-15 medium supplemented with 1× penicillin/streptomycin and 10% FBS
12. Pre-digestion mix: Leibovitz’s L-15 medium supplemented with 1× penicillin/streptomycin
13. Digestion mix: Pre-digestion mix supplemented with 2250 units/mL of trypsin and 3 mg/mL Collagenase A. Prepare fresh the day of dissociation, filter sterilise and store on ice until use. We recommend max. two pairs of 4^th^ and 5^th^ mammary glands (from two mice) per 10 mL of digestion mix.
14. Fibroblast clearing media: DMEM high glucose (subheading 2.2, item 3)
15. Mammary cell culture media: DMEM/F12 (subheading 2.2, item 2) supplemented with 1× penicillin/streptomycin and 10% FBS
16. Dissection tools: sharp surgical scissors and fine forceps
17. Ice
18. Ethanol (70%)
19. gentleMACS™ Dissociator
20. gentleMACS™ C Tubes
21. Centrifuge tube (15 mL)
22. Tissue culture flask (75 cm^2^)
23. Optical grade (0.17 mm, #1.5) glass bottom sterile cell culture dishes
24. Physiological salt solution with 1.8 mM Ca^2+^ (PSS-Ca^2+^, see subheading 2.1, item 28)
25. *Optional*, CellTracker Red CMTPX
26. *Optional*, CellTracker Red CMTPX working solution (subheading 2.1, item 33)
27. Epifluorescence or confocal microscope
28. *Optional*, pharmacological agents to stimulate Ca^2+^ events in cells, for example 2× oxytocin solution (see subheading 2.1, item 31)
29. Humidified cell culture incubator (37°C, 5% CO_2_)
30. Biosafety cabinet

## 3. Methods

### 3.1. Mammary gland *ex vivo* Ca^2+^ imaging

#### 3.1.1 Mammary tissue preparation for *ex vivo* imaging

*When possible, prepare the microscope prior to euthanising the mouse to maximise imaging time with viable tissue*.

1. Euthanise animals according to local regulations. We employ different methods of euthanasia for embryos, neonates, and pregnant or lactating mice (**Note 5**). Start a laboratory timer to monitor time-post-cull (an important consideration for tissue viability).
2. Dissect the mammary glands as illustrated in **Figure 4a** and described in detail in Stewart & Davis [4] (**Note 6**).
3. Place the mammary gland in a large petri dish (**Figure 4b**) and cut the gland into 3-5 mm^3^ pieces without delay (**Figure 4c**, **Note 7 and 8**).
4. Place approx. 15-20 tissue pieces in a 50 mL centrifuge tube containing 10-15 mL of pre-warmed live imaging media (subheading 2.1, item 29), as shown in **Figure 4d**. *Optional*: incubate some tissue pieces in CellTracker Red working solution (subheading 2.1, item 33) for 20 minutes (**Note 9**). Loosen the lid of the centrifuge tube to allow for some gas exchange.
5. Store tissue pieces in the humidified incubator at 37°C and 5% CO_2_. Tissue should ideally be used within 4-5 hours post-dissection.
6. Place a mammary tissue piece in the centre of an imaging dish (**Figure 4e**) and anchor it in place using a piece of net and a washer as shown in **Figure 4f and g**.
7. Add 200 µL of PSS-Ca^2+^ (subheading 2.1, item 28) to the tissue piece and immediately proceed to the confocal microscope (**Notes 10 and 11**).

**Figure 4:**
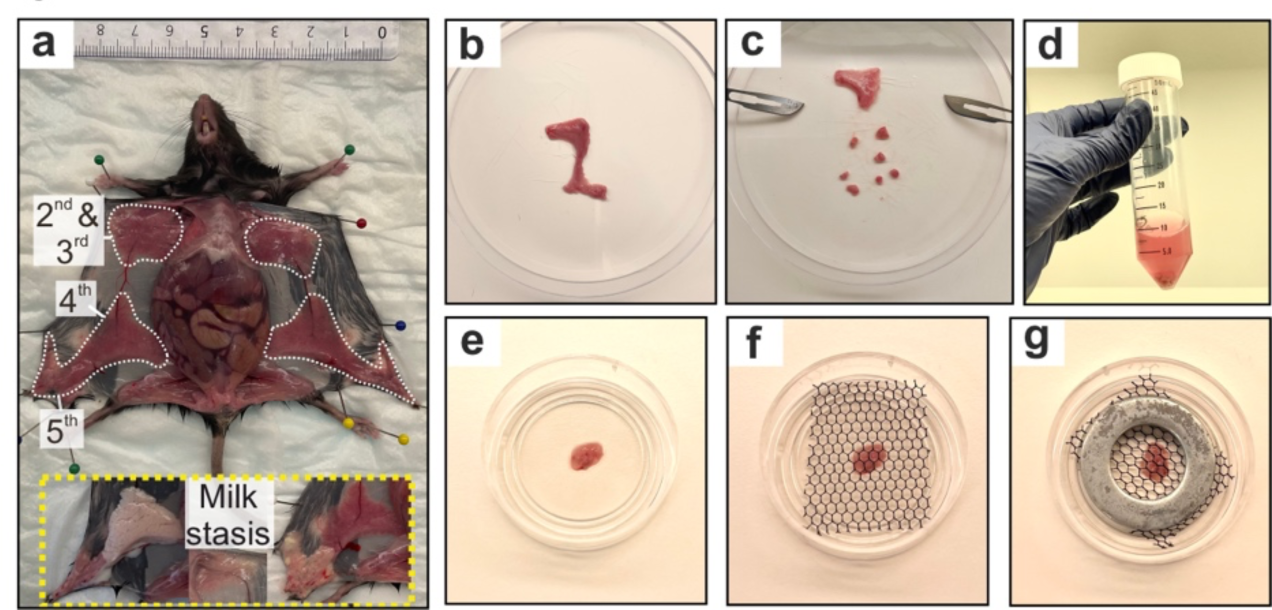
Mammary tissue preparation for *ex vivo* imaging. a. Mouse thoracic (2^nd^ and 3^rd^, or “upper”), abdominal (4^th^), and inguinal (5^th^) mammary glands during lactation. Inset (yellow dotted box) shows mammary glands presenting with milk stasis that are thus generally considered not suitable for live imaging. b. 4^th^ and 5^th^ mammary glands, cut into pieces (c) and incubated in live imaging media (d). e. Preparation of a mammary gland piece for live imaging using a net (f) and a washer (g) to minimise tissue displacement upon liquid addition.

#### 3.1.2. Ex vivo imaging of mammary tissue

*Data presented in this chapter have been obtained using either a resonant scanning Olympus FV3000 confocal microscope or a resonant scanning Leica STELLARIS 8 confocal microscope. Imaging settings will depend on the confocal system available in your laboratory and thus we provide only very general guidelines in this section*.

1. Prepare the imaging system. Imaging settings routinely used by our laboratory include:
  a. Use of long working distance water or silicone immersion objectives (25-30×).
  b. Imaging GCaMP6f and CellTracker (if used) on the same track to improve acquisition speed.
  c. Use of a resonant scanner and bidirectional scanning to improve acquisition speed.
2. Focus on the sample and identify a region of interest (ROI) for imaging (**Note 12**).
3. Define the beginning and end of the volume/stack. Typically, we acquire 30-40 µm volumes (approx. 7-10× 3-5 µm optical slices) in approximately 3.5 s.
4. Image for 2-5 minutes prior to agonist addition to establish a baseline.
5. Add 200 µL of 2× oxytocin (subheading 2.1, item 31) to the dish containing 200 µL of PSS-Ca^2+^ and continue imaging for 20-25 minutes.
6. After data acquisition is complete, images can be visualised using the Bio-Formats plugin (ImageJ) [41, 42]

### 3.2. Primary mammary cell isolation and *in vitro* Ca^2+^ imaging

#### 3.2.1 Mammary epithelial cell isolation

*Expected time to complete procedure:* 8-10 hours, of which 30 minutes (step 3), 1 hour 15 minutes (step 10) and 4-6 hours (step 15) are incubation times.

1. Euthanise animals according to local regulations. Dissect abdominal and inguinal mammary glands (**Figure 5a**, **Note 13**). Briefly dip tissue in ethanol (70%) to help maintain sterility (particularly if dissecting on the bench).
2. Carefully remove the lymph nodes and roughly dice mammary tissue (**Figure 5b**).
3. Transfer tissue to a gentleMACS™ C Tube containing 10 mL of digestion mix (subheading 2.2, item 13). Place the gentleMACS™ C Tube in the gentleMACS™ Dissociator and run the pre-set programme 37C_m_LDK_1 (**Figure 5c, Note 14**).
4. From this point forward, all work is completed in a biosafety cabinet using aseptic technique. At the completion of the gentleMACS^TM^ programme, transfer the contents of the gentleMACS™ C Tube to a 15 mL centrifuge tube. Centrifuge the sample for 5 minutes at 1500 rpm.
5. Following centrifugation, transfer the fat layer (**Figure 5d**) and supernatant to a fresh 15 mL tube (tube 2). Resuspend the contents via pipetting and centrifuge tube 2 for a further 5 minutes at 1500 rpm.
6. Meanwhile, resuspend the pellet from the original tube with 1 mL dissociation media (subheading 2.2, item 11) and transfer to a new 15 mL tube (tube 3). Wash the original tube with 2 mL of dissociation media and transfer to tube 3 to collect any remaining cells.
7. Following centrifugation of tube 2, remove the supernatant and resuspend the small pellet in 1 mL dissociation media. Transfer to tube 3 and centrifuge for 5 minutes at 1500 rpm (**Figure 5e**, **Note 15**).
8. Remove the supernatant and resuspend the pellet in 5 mL of red blood cell lysis buffer. Incubate at room temperature for 5 minutes (**Figure 5f**).
9. Add 10 mL of dissociation media and centrifuge at for 5 minutes at 1500 rpm (**Note 16**).
10. Remove the supernatant and resuspend the cells thoroughly but gently in 1 mL of fibroblast clearing media (subheading 2.2, item 14) with a P1000 pipette. Transfer to a 75 cm^2^ tissue culture flask. Wash the 15 mL tube with 12 mL of fibroblast clearing media to collect any remaining cells and transfer to the 75 cm^2^ tissue culture flask. Gently shake the flask in a horizontal plane and incubate in a humidified incubator for 1 hour and 15 minutes at 37°C and 5% CO_2_ (**Figure 5g**).
11. After the incubation, most of the fibroblasts will attach to the tissue culture plastic whilst epithelial cells will not. Shake the flask in a horizontal plane with moderate vigour and transfer media to a new 15 mL tube. Rinse the tissue culture flask with another 13 mL of dissociation media and transfer to another 15 mL tube. Centrifuge both cell suspensions for 5 minutes at 1500 rpm (**Figure 5h**).
12. Remove the supernatants and combine pellets by resuspending mammary cell culture media (subheading 2.2, item 15).
13. Count the cell suspension and adjust solution to a concentration of 5 million cells/mL.
14. Prepare imaging plates by pipetting approx. 5 µL droplets of cell suspension in the centre of the dish. To maintain humidity within the dish, fill the rim of the dish with droplets of cell-free complete media (**Figure 5i**). Plating the cells in this way maximises the amount of imaging dishes that can be created and assists in minimising adhesion time for the cells.
15. Incubate the plated cells at 37°C, 5% CO_2_ in a humidified incubator for 4-6 hours until cells have adhered. At the end of incubation, gently wash the dish with 500 µL mammary cell culture media to remove any material that has not adhered. Replenish the dish with mammary cell culture media.
16. Isolated cells should be imaged with 24 hours of dissection.

**Figure 5:**
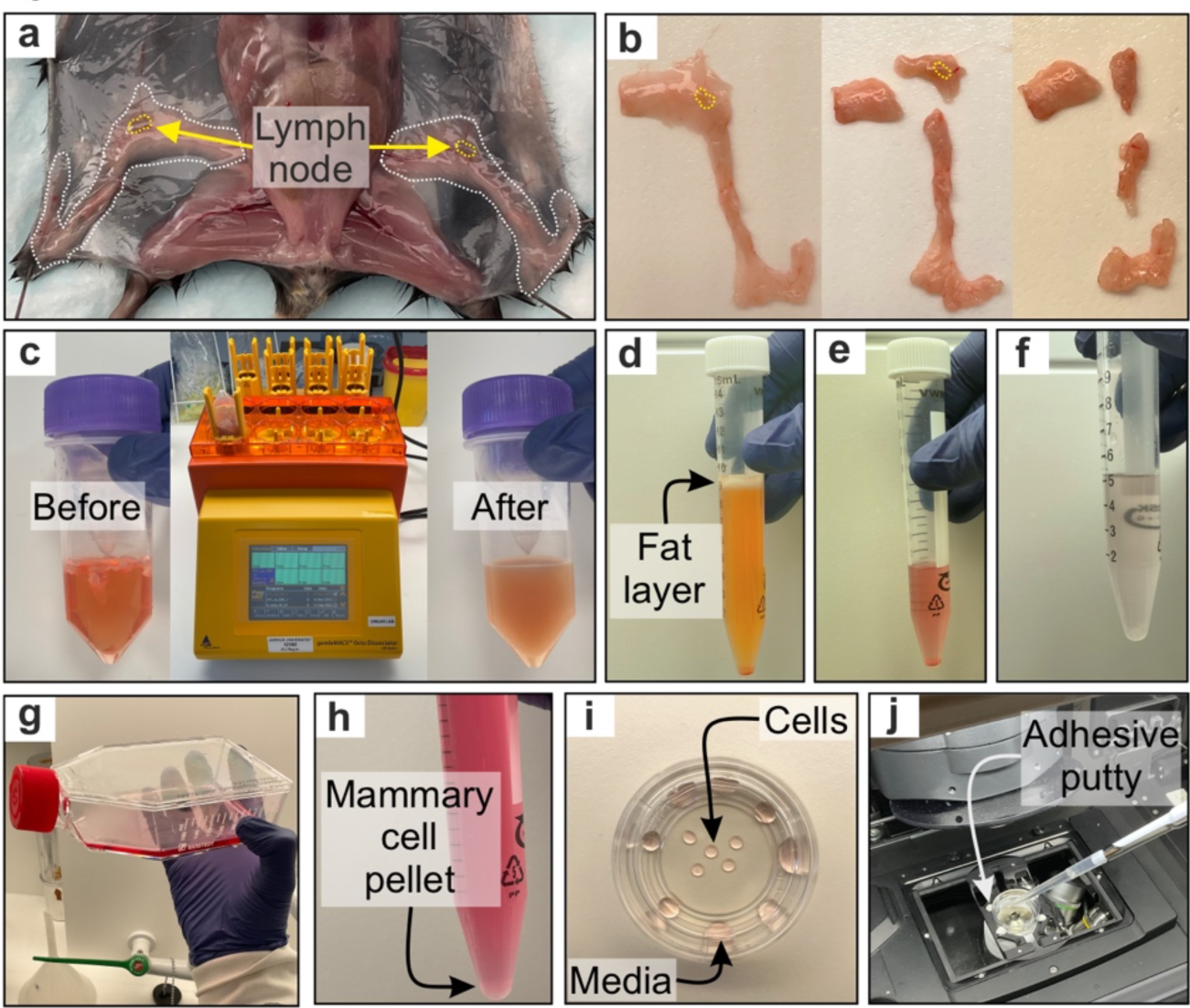
Mammary tissue preparation for *in vitro* imaging. a. Nulliparous mouse abdominal (4^th^), and inguinal (5^th^) mammary glands (white dashed outline) with lymph nodes identified by yellow arrow and outline. b. Dissected mammary glands should have lymph nodes identified (yellow dashed line, panel 1) and removed (panel 2) prior to dicing roughly (panel 3). c. Diced mammary glands are dissociated via gentleMACS™ Dissociator. d. Dissociated mammary tissue post-centrifugation, prior to removal of fat layer and supernatant, and after removal (e). f. Following incubation in red blood cell lysis buffer, the cell suspension becomes colourless. g. Fibroblasts are removed via short incubation in a cell culture flask. h. Cell pellet at the end of isolation protocol should be clearly visible at the bottom of a 15 mL centrifuge tube. i. Following isolation, mammary cells are counted and plated in imaging dishes via the droplet approach. j. An example of liquid addition onto an imaging plate mounted on a confocal microscope. Adhesive putty placed gently around the imaging plate can minimise movement artifacts due to liquid addition.

#### 3.2.2 *In vitro* imaging of isolated mammary epithelial cells

*Data presented here have been obtained using either an Olympus FV3000 confocal microscope or a Leica STELLARIS 8 confocal microscope, however confocal capabilities are not a requirement for imaging* in vitro *2D samples*.

7. *Optional*: Load the cells with CellTracker Red by incubating the plates in CellTracker working solution (subheading 2.2, item 26) for at least 30 minutes at 37°C and 5% CO_2_. Following incubation, replace CellTracker loading media with mammary cell culture media (subheading 2.2, item 15) until ready to image.
8. Prepare the imaging system. Imaging settings routinely used by our laboratory include:

a. Imaging via water or silicone objectives.
b. Imaging GCaMP6f and CellTracker (if used) on the same track to maximise acquisition speed.
c. Use of a resonant scanner and bidirectional scanning to improve acquisition speed.
d. A small volume (*z* stack) of approximately 4.5 µm, (3× 1.5 µm optical slices) in approx. 1 second.
e. Adaptive Focus Control (or an equivalent *z*-drift compensator).
9. Replace the media on one imaging dish with 1mL PSS-Ca^2+^ (**Note 10**).
10. Transfer the sample to the imaging system. To reduce the likelihood of the imaging plate moving during liquid addition, a small amount of adhesive putty can be gently placed around the plate on the stage (**Figure 5j**).
11. Identify a region of interest and commence imaging. After approximately 40 seconds, gently add 1 mL of 2× agonist solution gently onto the imaging plate (**Figure 5j**).

### 4.0 Notes

1. Tamoxifen has a dose dependent effect on mammary gland development [4]. Tamoxifen dose and method of administration should be optimised in tamoxifen-dependent models to suit the experiment. Pregnant mice that have previously been administered tamoxifen should be monitored for dystocia around the time of littering.
2. Most GECIs have low fluorescence at resting cytosolic Ca^2+^ concentrations. CellTracker Red is useful for the identification of ROIs in tissue pieces. If imaged, the red channel can also be used to visualise the nature and extent of alveolar warping upon oxytocin stimulation. Other dyes in the CellTracker family (or other live cell stains) could be considered, however, the best results in our laboratory have been obtained using CellTracker Red CMTPX. This dye preferentially stains luminal cells (see Fig S1A, Ref [16]).
3. This protocol is suitable for mice of any age; however, mice < 8-wks old (where ductal morphogenesis is incomplete) may have a lower yield of mammary epithelial cells. Samples isolated from mice > 20-wks old may have greater variability in yield and purity.
4. This protocol has been optimised for isolation of cells from nulliparous mammary tissue. In our experience, cell isolation during lactation is problematic due to epithelial cell death. A method for isolating basal cells during gestation is outlined in Ref [16] and includes use of nano-patterned dishes for optimal basal cell attachment.
5. We euthanise pregnant and lactating mice by CO_2_ inhalation for 10 minutes or 5 minutes, respectively. Embryos from pregnant mice are immediately removed from the mother’s uterus and decapitated using sharp scissors. Due to their immature lung development, we typically euthanise 10-12-day old neonates by intraperitoneal injection of a lidocaine-pentobarbital mixture, followed by swift decapitation with sharp scissors.
6. For lactation studies, experiments are ideally performed on lactation days 10-12 (peak lactation) on mice nursing 6 or more pups. When litter size < 6, some mammary glands may not be frequently or adequately suckled and may appear white, blotchy or streaky (**Figure 4a, milk stasis insert**). This can signify partial involution (or in some cases mastitis) and the glands should generally be avoided.
7. We often favour the 5^th^ and upper mammary glands for pharmacological studies. These glands are thinner and thus allow faster diffusion of agonists/antagonists through the tissue to the ROI. Nevertheless, it is also reasonable and appropriate to use the 4^th^ mammary gland for *ex vivo* imaging.
8. To avoid ripping-, tearing- or crushing-like damage at the edge of tissue pieces, we prefer to dice mammary tissue between two #20 scalpel blades pulled swiftly in opposite directions. In our hands, this method causes less damage than cutting with dissection scissors or by pressing the gland under the force of a single blade. Do not allow the gland to dry out during cutting.
9. Tissue can remain incubated in CellTracker Red working solution for several hours.
10. Samples should not remain in PSS-Ca^2+^ for more than 10 minutes prior to imaging.
11. Remove as much culture medium as possible from the tissue piece before adding the PSS-Ca^2+^.
12. CellTracker Red can help to identify mammary alveoli down the eyepiece. Ideally alveoli within the field of view should be close to the edge of the tissue piece (approx. 2-3 alveolar diameters from the edge), to ensure that the agonist consistently diffuses to the ROI within several minutes of application. Damaged regions (typically characterised by high, uniform resting GCaMP6f fluorescence or a broken/damaged appearance) should be avoided. Alveoli on the outermost edge of the tissue are more subject to motion artifacts when the agonist is applied, particularly when using oxytocin, which causes basal cells to contract and alveolar structures to shift in *x*, *y* and *z* planes. This can lead to the ROI moving out of the field of view or out-of-focus during acquisition. Even for the most experienced investigator, motion artifacts are a major challenge during imaging.
13. Thoracic mammary glands are less suitable for this protocol due to the propensity of these glands to be contaminated with surrounding skeletal muscle.
14. This is a pre-set, proprietary program the details of which have not been disclosed to us by the distributor. If adopting a different system, the program will need to be optimised for mammary tissue. The program involves heating to 37 °C for approximately 30 minutes with agitation.
15. Steps 5-7 enable the recovery of maximal material, but could be skipped to save time if more than enough material are recovered at the end of step 4.
16. If pellet remains red following red blood cell lysis, steps 8 and 9 could be repeated.

## Acknowledgments

This work was supported by the Novo Nordisk Foundation (NNF20OC009705), the Carlsberg Foundation (CF21-0170) and the National Health and Medical Research Council of Australia (2003832). We thank Drs Teneale Stewart and Bethan Lloyd-Lewis, as well as Mr Alexander Stevenson, for their help developing and optimising these protocols. We thank Ms Trine Lund Ruus and Dr Amita Gautam Ghadge for their animal and laboratory support. We would also like to thank the TRI Microscopy Core Facility (University of Queensland), the Katharina Gaus Light Microscopy Facility (University of New South Wales) and the Bioimaging Core Facility (Aarhus University). Figures were created in-part with BioRender.com.

